# On the optimal use of isotherm models for the characterization of biosorption of lead onto algae

**DOI:** 10.1101/035774

**Authors:** F. Brouers, Tariq J. Al-Musawi

## Abstract

For the first time we apply a new method based on the mathematical derivation of some known isotherm from the Burr function which describes many birth-death (sorption-desorption) phenomena in ecology and economy. Therefore, in this study the experimental isotherm data of biosorption of Pb(II) onto algae was modeled to Langmuir, Hill-Sips, Brouers–Sotolongo, Brouers-Gaspard, and Redlich-Peterson isotherm models. The parameters of each model were determined by non linear fitting algorithms using Mathematica program. The maximum Pb(II) removal rate increased with the increase of temperature and reached the maximum value (98%) at the temperature of 40 °C. The results showed that the Hill-Sips and the Brouers–Sotolongo isotherms were definitely the most suitable models to satisfactorily describe biosorption of Pb(II) on the algal biomass. In addition, as these two models gave very close results, the use of an intermediate one the Brouers-Gaspard isotherm model could also describe the sorption in most cases. High coefficient of determination values were obtained by using non linear methods and these findings are contrary to most works in this field that use linearization methods. Further, this study showed that a complete set of data is necessary to have a good representation of the isotherm and using only coefficient of determination is not always an adequate tool to compare the goodness of the non linear fit of an isotherm models.

## 1. Introduction

Biosorption is a process which utilizes inexpensive dead biomass to sequester toxic pollutants (**Kratochvil and Volesky, 1998**; **Davis et al., 2003**, **Gupta et al., 2012a**). Biosorption is proven to be quite effective at removing metals ions from contaminated solutions in a low cost and environment-friendly manner. Adsorbent comes under the following categories: algae; fungi; grass, agriculture waste, and nanomaterials (**Sulaymon et al., 2013a; Holan and Volesky, 1995; Sulaymon et al., 2014**; **Mittal et al., 2010a**; **Mittal et al., 2009a**; **Saleh and Gupta, 2012a**; **Saleh and Gupta, 2012b**). These materials were used to remove pollutants in a batch and/or continuous systems (**Gupta et al., 2011a**, **Mittal et al., 2010b**; **Gupta and Nayak, 2012**; **Khan et al., 2010; Karthikeyan et al., 2012**). The term algae refers to a large and diverse assemblage of organisms that contain chlorophyll and carry out oxygenic photosynthesis (**Davis et al., 2003**). Algae are one of the most abundant and highly available natural resources in tropical ecosystems. Cyanophyta (blue–green algae), Chlorophyta (green algae), Rhodophyta (red algae), and Phaeophyta (brown algae) are the divisions of the largest visible algae. Many studies concluded that the algal biomass is a very promising material to be used as biosorbent to remove various kinds of pollutants from contaminated water and wastewaters (**Sulaymon et al., 2013b; Romera et al., 2007; Altenor, et al., 2012)**. This can be attributed that the algal biomass has many negative charge active groups on its surface cell wall such as hydroxyl, carboxyl, amino, sulfhydryl, and sulfonate (**Davis et al., 2003**).

The term isotherm is used for describing the retention of a substance on a solid at a constant temperature. The isotherm study is a major tool to predict the efficiency of a sorbent to remove a given pollutant from polluted water (**Ncibi et al., 2008; Gupta et al., 2011b**; **Jain et al., 2003**). Analysis of isotherm data is an interesting mathematical approach for describing sorption isotherms at a constant temperature for water and wastewater treatment applications and to predict the overall sorption behavior under different operating conditions (**Gupta et al., 2012b; Mittal et al., 2009b**). Hence, a large number of studies have been done to develop a mathematical isotherm models and to verify their suitability for describing the biosorption of heavy metals such as Pb^2+^; Cd^2+^, and Cr^3+^ by as well as to understand the sorption isotherm phenomenon between the biomass surface and the metal molecules. Most of these models are empirical and bring little information on the physicochemical processes responsible for the particular shape of the isotherm curves. **Brouers (2014a)** has shown that some of the most empirical models that can be used to describe the isotherm data were approximations of a generalized Brouers–Sotolongo model. This model is derived from the Burr-Madalla distribution which is itself solution of a birth and death differential equation which can describe sorption-desorption mechanisms (**Burr, 1942; Maddala, 1996**). The author concluded that only the Langmuir, the Hill-Sips and the Brouers–Sotolongo isotherms were genuine statistical functions and had the correct asymptotic limits for low and high concentrations. In addition, the author pointed out that a statistical analysis had a meaning only if the experimental data were complete until the saturation of the sorption. Furthermore, Ion-exchange is an important concept in biosorption mechanisms using algae, because it explains many of the observations made during heavy metal uptake experiments. In this case, the ion exchange reaction type occurs between light metals already bound to the algae and other metals present in the aqueous solution (**Naja and Volesky, 2006)**. The ion-exchange model is certainly a better representation of the biosorption process using algae since it reflects the fact that most algal biomass is either protonated or contains light metal ions such as K+, Na+ and Mg^2^+, which are released upon binding of a heavy metal cation. It should be pointed out that the ion exchange model does not explicitly identify the binding mechanism (**Davis et al., 2003)**. Hence, verification of the various isotherm models is a growing area of study.

Despite the fact that linear regression is still frequently used (**Sulaymon et al., 2010**), non-linear analysis of isotherm data is an interesting and useful mathematical approach for describing biosorption isotherms for water and wastewater treatment applications and to predict the overall sorption behaviour under different operating conditions (**Ho, 2006**). Indeed, the linearization of non linear isotherm equations has several disadvantages. This process implicitly alters their error structure and may also violate the error variance and normality assumptions of standard least squares (**Ratkowsky, 1990**). In general, non linear regression gives a more accurate determination of model parameters than linear regression method. In recent years, several error analysis methods, such as the coefficient of determination (R^2^), the sum of the errors squared, a hybrid error function, Marquardt’s percent standard deviation, the average relative error and the sum of absolute errors, have been used to determine the best-fitting isotherm (**Ho et al., 2002; Allen et al., 2003**). **Ho (2004)** pointed out that using only R^2^ of linear regression analysis is not an appropriate tool to evaluate the goodness of the fit of an isotherm model. As different forms of the equations affected R^2^ values more significantly during the linear analysis, the non-linear analysis might be a method of avoiding such errors (**Ho and Wang, 2004**; **Kumar and Sivanesan, 2005**). Additionally, **Brouers (2014b)** showed that the use of nonlinear programs allows an easy and precise fitting of the experimental data with the theoretical models.

The aim of this paper is to apply and demonstrate the interest of new methodology introduced by **Brouers (2013, 2014a)** on the isotherm data for the biosorption of Pb(II) in aqueous solution using algae. The non linear method of five isotherm model, Langmuir, Hill-Sips, Brouers–Sotolongo, Brouers-Gaspard, and Redlich-Peterson models, were compared with the experimental data. A trial-and-error procedure was used for the non-linear analysis method using the Mathematica 9 program and based on the Burr-Madalla distribution function introduced by **Burr (1942)**.

## 2. Materials and Methods

### 2.1 Materials

Fresh mixture of green and blue–green algae was used in this study as a biosorbent. This material was collected from the artificial irrigation canal in Baghdad University, Iraq. It was mainly a mixture of three species of algae. Blue–green Oscillatoria princeps alga was the highest percentage (88%), green Spirogyra aequinoctialis alga was (9%), and blue-green Oscillatoria subbrevis alga (3%). And to make it user friendly the collected algae were not separated. The foreign matters were removed manually from the collected algae, then rinsed with tap and distilled water to remove of dirt, sands, and external salts. Afterward, the washed algae were kept in air for removing water and dried at an oven temperature of 65 °C for 48 h. The dried biomass were roughly chopped, grounded into powder, sieved, and kept in air-tight polyethylene container at room temperature. An average size of 0.54 mm was used for experiments with required amounts. A sample of algal biomass was analyzed for the physico-chemical properties and the results were listed in Table (1). The surface area was determined from N_2_ adsorption isotherm using a Micrometrics Nano Porosity System. Point of zero charge (pH_pzc_) of used biosorbents was determined by conventional method (**Kalhori et al., 2013**). Briefly, initial pH values of 50 mL of 0.1 M NaCl were adjusted in a pH range of 2 to 12 using 1 M NaOH or H_2_SO_4_ solutions. Afterward, a 0.5 g of algal biomass was added to each solution and the obtained suspensions were shaken for 48 h. The algal biomass was filtered and the final pH of the solution was measured. The value of pH_pzc_ of algal biomass was found from the intersection of the curve of final vs. initial pH. The micropore volume was evaluated by the t-plot method. Total exchange capacity was measured using atomic absorption spectrophotometer (type: SHIMADZU, AAS 7200, Japan).

**Table 1.**
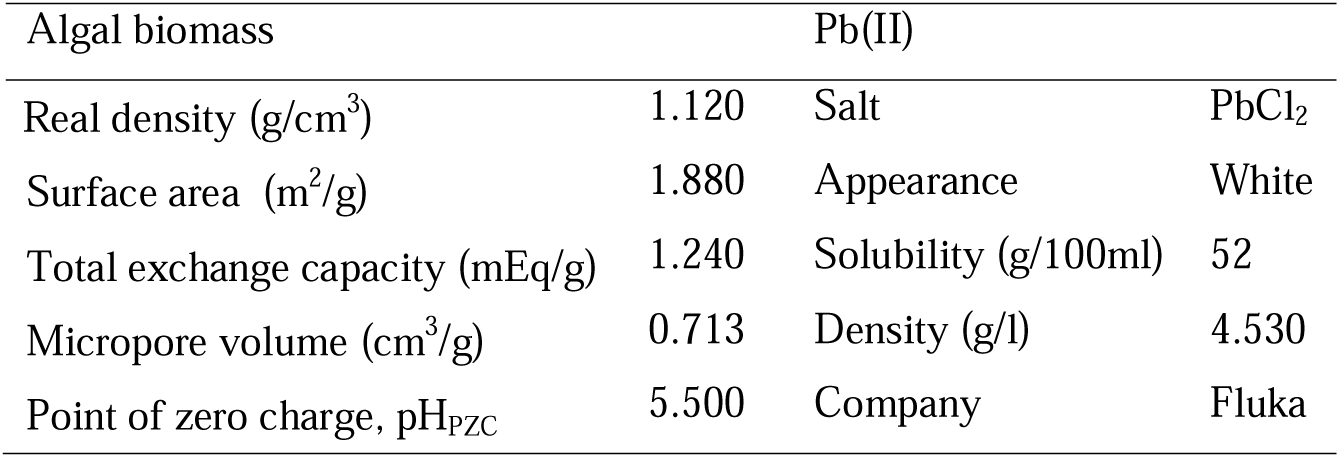
Properties of the algal biomass and Pb(II) salt used in this study

All the chemicals used in this work are analytical grade reagents with deionized water used for solutions preparation. Stock solution (1000 mg/l) of Pb(II) was prepared by dissolving the appropriate weight of lead chloride in distilled water and kept in glass container at room temperature. The desired concentrations were prepared by diluting the stock solution in accurate proportions to different initial concentrations. The initial pH of the working solutions was adjusted by addition of 1 mol/l NaOH or HCl using a pH meter (WTW, inoLab 720, Germany). All the glassware used for dilution, storage and experimentation were cleaned with detergent, thoroughly rinsed with tap water, soaked overnight in a 20% HNO3 solution and finally rinsed with distilled water before use. Table (1) listed the properties of Pb(II) salt used in this study.

### 2.2 Methods

#### 2.2.1 Experiments

Isotherm experiments were carried out in 250 ml stoppered conical flasks containing 0.05, 0.1, 0.3, 0.5, 0.8, 1, 2, and 3 g of algal biomass and 100 ml of Pb(II) solution. These experiments were performed at the same initial concentration (50 mg/l) for Pb(II) and repeated for temperature range from 10 to 40 °C. According to the previous studies, the optimum pH for Pb(II) removal using algae is in the range between pH 2 and pH 4 (**Sulaymon et al., 2013a**; **Davis et al., 2003**). In addition, several authors showed that further increases in temperature (above 40 °C) lead to a decrease the percentage removal. This may be attributed to an increase in the relative desorption of the metal from the solid phase to the liquid phase, deactivation of the biosorbent surface, destruction of active sites on the biosorbent surface **(Saleem, et al., 2007; Meena, et al., 2005**. Hence, in this study all the isotherm experiments were conducted at temperature below 40 °C and the pH solution was adjusted to the 3. The flasks were placed in a shaker (Edmund Buhler, 7400 Tubingen Shaker-SM 25) with constant shaking at 200 rpm for 4 h. After equilibrium condition, the sorbent was separated from aqueous solution by using filter paper (WHATMAN, No.42, diameter 7 cm). The Pb(II) concentrations in both initial and withdrawn samples were determined using atomic absorption spectrophotometer (type: SHIMADZU, AAS 7200, Japan). Each sample was measured thrice and the results were given as the average value. Biosorption capacity q(x) at equilibrium conditions and the percentage of removal (%) were calculated using eq(1) and eq(2), respectively.

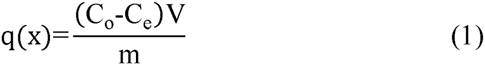

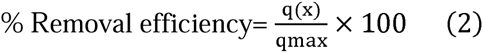

where, q(x) and q_max_ are the equilibrium and maximum biosorption capacity of Pb(II) (mg/g), respectively; C_o_ and C_e_ are the initial and equilibrium Pb(II) concentrations in the aqueous solution (mg/l), respectively; V is the volume of used solution (l); and m is the mass of used adsorbent (g).

#### 2.2.2 Isotherm models

There are several isotherm models with two or more than two parameters available for analyzing the experimental parameters. The parameters of each model often provide insights into the sorption mechanism, the surface properties and the affinity of the sorbent (**Yu and Neretnieks, 1990**). In the present study, five different isotherm models were tested under different adsorption temperatures. These models are: Langmuir, Sips, Brouers–Sotolongo, Brouers-Gaspard, and Redlich-Peterson models were chosen to fit the experimental data and were derived from the General Brouers–Sotolongo equation as will be shown below (**Brouers, 2014a)**. The non linear regression analysis using Mathematica program, version 9, was used for direct determination of each model parameters. In this case, a trial-and-error procedure, which is applicable to computer operation, was developed to determine the isotherm parameters using an optimization routine to maximize the coefficient of determination (R^2^) between the experimental data and isotherms. The applicability and suitability of the isotherm equation to the equilibrium data were compared by judging the values of the R^2^ values.

The General Brouers–Sotolongo (GBS) isotherm is represented by the following equation (**Brouers, 2014a**):

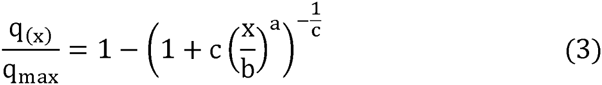

Where q_max_ is the maximum sorbed quantity (mg/g), x is the residual metal concentration in solution at equilibrium (mg/l), b is the isotherm constant (mg/l), a is a coefficient dependent on the fractal nature of the system and this exponent is a measure of the width of the adsorption energy distribution and energy heterogeneity of the sorbent surface (**Ahmad et al., 2014**), and c is the coefficient related to its cluster organization (**Stanislavsky and Weron, 2013**).

If c=1 and a=1, GBS reduces to the Langmuir model (eq. (4)). Langmuir model is the best known and most often used isotherm model for the sorption of a solute from a liquid solution. This model is based on the assumption that sorption sites are identical and energetically equivalent, and that the solute is immobilized under the form of monolayer coverage (**Langmuir, 1918**).

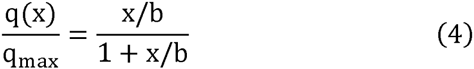

**Hill-Sips** (HS) isotherm model (eq.(5)) can be derived from GBS when c=1. Hill-Sips isotherm model is a combined form of Langmuir and Freundlich expressions deduced for predicting the heterogeneous adsorption systems and circumvent the limitation of the rising adsorbate concentration associated with Freundlich isotherm model (**Ahmad et al., 2014**). At high adsorbate concentration, it predicts monolayer adsorption characteristics of Langmuir, while in low adsorbate concentration, it reduces to Freundlich isotherm (**Sips, 1948**).

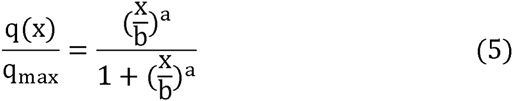

For c=0, one recovers the Brouers–Sotolongo (BS) isotherm equation (eq. (6)). Brouers–Sotolongo (BS), which is basically a deformed Weibull exponential equation, was specifically developed for the complex and high heterogeneous systems and the fact that this model considers a pattern of sorption energy distribution makes it suitable to describe adsorption phenomena involving sorbing materials even if they show different chemical and structural characteristics (**Altenor, et al., 2012**). The surface of the biosorbent is assumed to be made of a finite number of patches of sites of different sorption energies (**Brouers et al., 2005**).

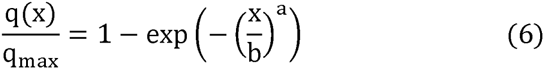

Indeed, it is difficult in many cases to choose between HS and BS isotherms, it is why it has been proposed to use in these cases an intermediate value for c (c=1/2) and introduce a new intermediate formula which is named Brouers-Gaspard (BG) isotherm equation (eq.(7)).

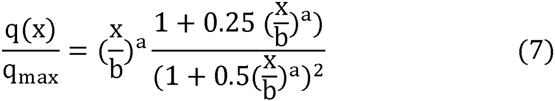

All these equations have good physical asymptotic behaviors:

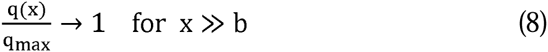

and

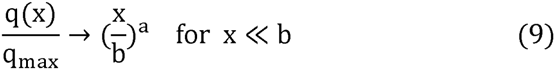

Another commonly used empirical isotherm is the Redlich-Peterson isotherm (RP) (eq. (10)). Redlich-Peterson isotherm model incorporates the features of the Langmuir and Freundlich models into a single equation and presents a general isotherm. This model can be applied to express the sorption process when dealing with certain pollutants at high concentration (**Quintelas, et al., 2008**).

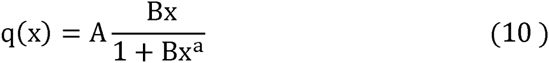

This equation does not have the physical correct asymptotic limit since it gives:

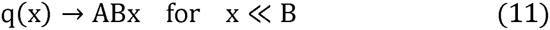

and

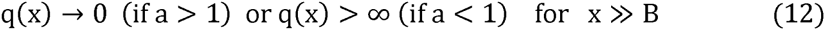

Where A and B are the Redlich-Peterson isotherm constants.

## 3. Results and discussion

The bisorption isotherm of Pb(II) onto algal biomass was investigated as a function of temperature and the results are depicted in Fig. 1 (a). The graph are plotted in the form of Pb(II) removal efficiency against algal biomass weight. These data may help in understanding the controlling mechanisms and quantifying the sorption properties of the sorbent, maximum sorption capacity and affinity of the sorbent for the target metal (**Langmuir, 1918**). For all experimental data shown in this figure, it can be seen that the removal efficiency was increased with the increase of algal biomass weight from 0.05 to 1 g, implying that the optimum amount of biosorbent is 1 g algal biomass/200 ml solution, then beyond 1 g algal biomass dose the removal efficiency reached a plateau which demonstrating the equilibrium state. The reason being that an increase in the biosorbent further increment in sorbent dose, the removal capacity was not increased possibly due to the aggregation of sorbent particles and low surface area. Moreover, Fig. 1 (a) shows that with an increase in temperature, the percentage of Pb(II) removal increases and the maximum removal efficiencies were obtained at 40 °C. This means that Pb(II) binding on active sites of the biosorbent becomes stronger at higher temperature and that the sorption process is endothermic. This can be attribute to the increase in temperature is knowing to increase the rate of diffusion of the adsorbate molecules across the external boundary layer and in internal pores of the adsorbent particles as a result of the reduced viscosity of the solution (**Bulut et al., 2012**). A similar trend was obtained by **Sulaymon et al., (2013b)**.

**Fig. 1.**
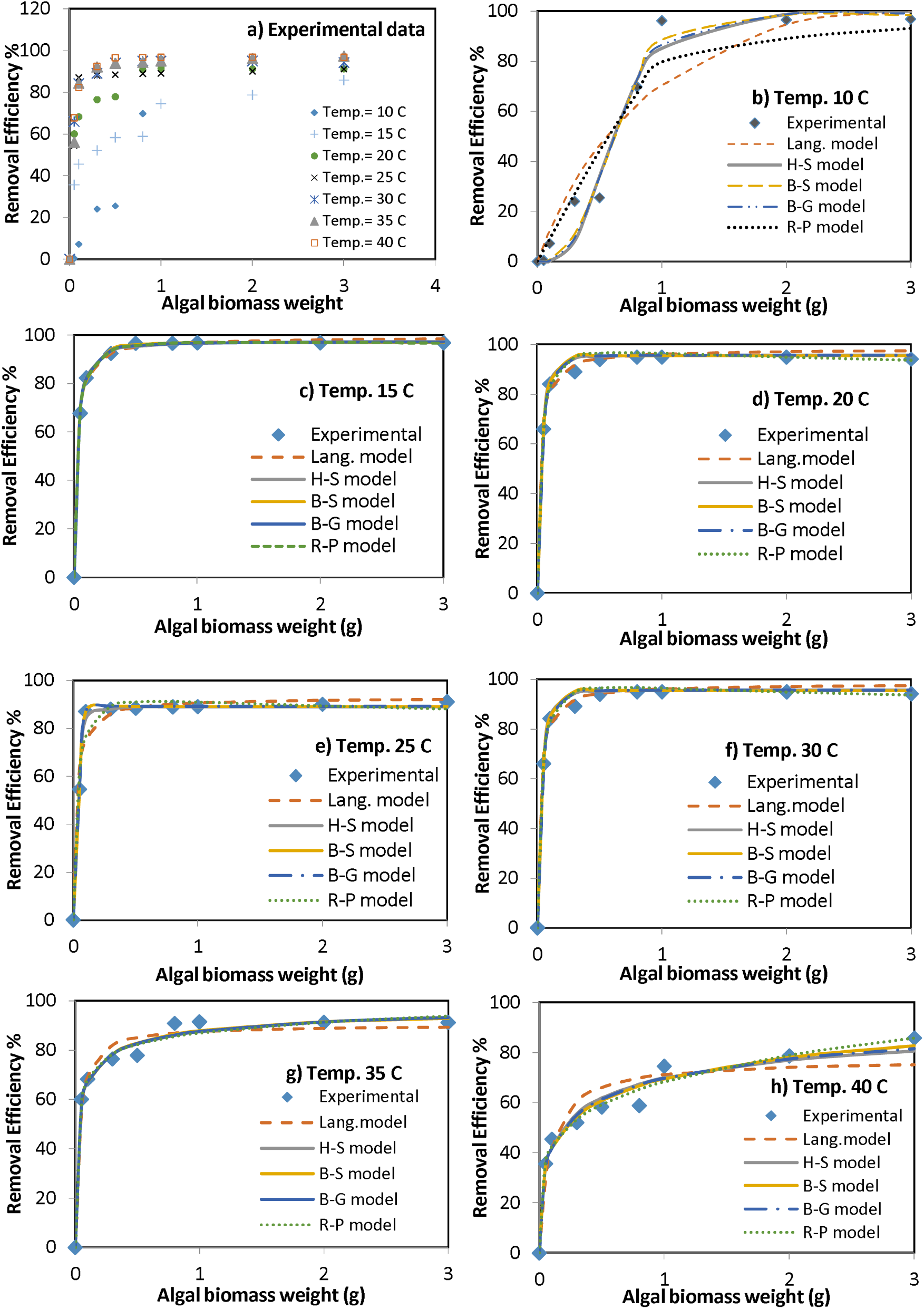
Experimental data of biosorption isotherm of Pb(II) ions at different temperatures using algal biomass and simulation of isotherm data with several models

The experimental isotherm data have been systematically modeled by the aforementioned isotherm models and the results are depicted in Fig. 1(b-h). The parameters of these models and the coefficient of determination (R^2^) values are listed in Tables 2. Based on the analysis of the estimated variance (R^2^ values), the B-S model gave a best fit of experimental data for all temperature values. The R^2^ values of Brouers–Sotolongo model were much closer to 1.0 than the values obtained with the other models. Such fitting tendency would refer to the presence of active sites with heterogeneous sorption interactions (**Ncibi, et al., 2008; Altenor et al., 2012**). In order to find out why this newly established Brouers–Sotolongo isotherm model was able to fit better the equilibrium data than the widely used models, a couple of points will be detailed hereafter. First of all, the B-S isotherm was an initiative to propose a new equation able to describe the adsorption process considering from the start that it is a complex system. Such trend would probably enlighten the energetic heterogeneity at the interface algal biomass. The heterogeneous surface stems from two sources known as geometrical and chemical ones. Geometrical heterogeneity is a result of differences in size and shape of pores, cracks and pits. Chemical heterogeneity is associated with different functional groups such as hydroxyl, amine, and carboxyl and aldehydes determining the apparent chemical character of an algal biomass surface (**Sulaymon et al., 2013a; Kratochvil and Volesky, 1998**). Indeed, under the used experimental conditions, the interactions involved in such bisorption case would be between lead molecules and algal biomass cell surfaces via different possible interactions mainly electrostatic attraction, ion exchange and complexation between lead and hydroxyl, amine, and carboxyl groups (**Davis et al., 2003)**. Thus, such tendency would lead to a highly heterogeneous sorption energy landscape, explaining therefore the good fit of the B-S isotherm model.

**Table 2.**
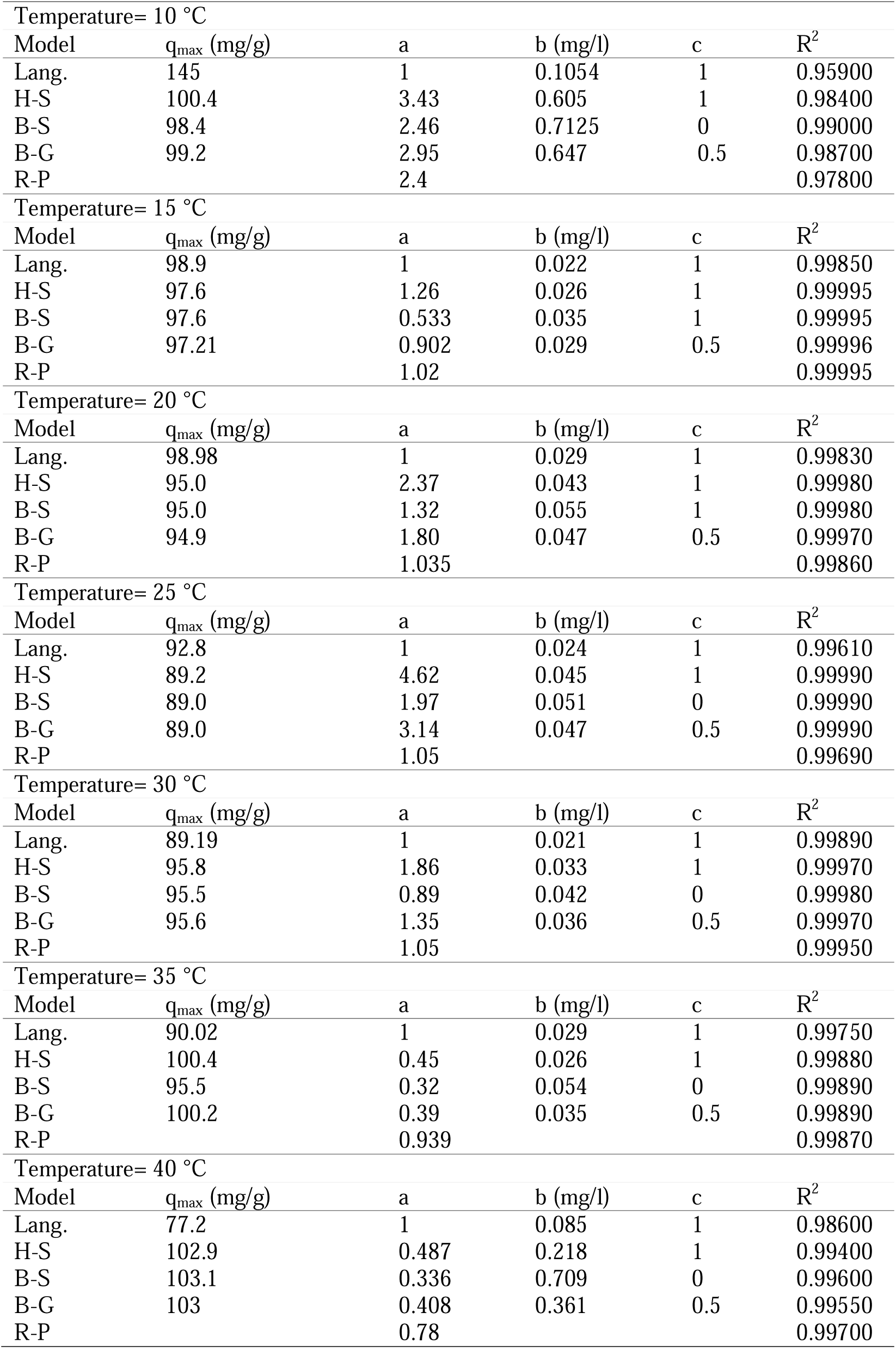
Isotherm modeling results related to the biosorption of Pb(II) onto algal biomass

| Temperature= 10 °C |  |  |  |  |  |
| --- | --- | --- | --- | --- | --- |
| Model | q<sub>max</sub> (mg/g) | a | b (mg/l) | c | R<sup>2</sup> |
| Lang. | 145 | 1 | 0.1054 | 1 | 0.95900 |
| H-S | 100.4 | 3.43 | 0.605 | 1 | 0.98400 |
| B-S | 98.4 | 2.46 | 0.7125 | 0 | 0.99000 |
| B-G | 99.2 | 2.95 | 0.647 | 0.5 | 0.98700 |
| R-P |  | 2.4 |  |  | 0.97800 |
| Temperature= 15 °C |  |  |  |  |  |
| Model | q<sub>max</sub> (mg/g) | a | b (mg/l) | c | R<sup>2</sup> |
| Lang. | 98.9 | 1 | 0.022 | 1 | 0.99850 |
| H-S | 97.6 | 1.26 | 0.026 | 1 | 0.99995 |
| B-S | 97.6 | 0.533 | 0.035 | 1 | 0.99995 |
| B-G | 97.21 | 0.902 | 0.029 | 0.5 | 0.99996 |
| R-P |  | 1.02 |  |  | 0.99995 |
| Temperature= 20 °C |  |  |  |  |  |
| Model | q<sub>max</sub> (mg/g) | a | b (mg/l) | c | R<sup>2</sup> |
| Lang. | 98.98 | 1 | 0.029 | 1 | 0.99830 |
| H-S | 95.0 | 2.37 | 0.043 | 1 | 0.99980 |
| B-S | 95.0 | 1.32 | 0.055 | 1 | 0.99980 |
| B-G | 94.9 | 1.80 | 0.047 | 0.5 | 0.99970 |
| R-P |  | 1.035 |  |  | 0.99860 |
| Temperature= 25 °C |  |  |  |  |  |
| Model | q<sub>max</sub> (mg/g) | a | b (mg/l) | c | R<sup>2</sup> |
| Lang. | 92.8 | 1 | 0.024 | 1 | 0.99610 |
| H-S | 89.2 | 4.62 | 0.045 | 1 | 0.99990 |
| B-S | 89.0 | 1.97 | 0.051 | 0 | 0.99990 |
| B-G | 89.0 | 3.14 | 0.047 | 0.5 | 0.99990 |
| R-P |  | 1.05 |  |  | 0.99690 |
| Temperature= 30 °C |  |  |  |  |  |
| Model | q<sub>max</sub> (mg/g) | a | b (mg/l) | c | R<sup>2</sup> |
| Lang. | 89.19 | 1 | 0.021 | 1 | 0.99890 |
| H-S | 95.8 | 1.86 | 0.033 | 1 | 0.99970 |
| B-S | 95.5 | 0.89 | 0.042 | 0 | 0.99980 |
| B-G | 95.6 | 1.35 | 0.036 | 0.5 | 0.99970 |
| R-P |  | 1.05 |  |  | 0.99950 |
| Temperature= 35 °C |  |  |  |  |  |
| Model | q<sub>max</sub> (mg/g) | a | b (mg/l) | c | R<sup>2</sup> |
| Lang. | 90.02 | 1 | 0.029 | 1 | 0.99750 |
| H-S | 100.4 | 0.45 | 0.026 | 1 | 0.99880 |
| B-S | 95.5 | 0.32 | 0.054 | 0 | 0.99890 |
| B-G | 100.2 | 0.39 | 0.035 | 0.5 | 0.99890 |
| R-P |  | 0.939 |  |  | 0.99870 |
| Temperature= 40 °C |  |  |  |  |  |
| Model | q<sub>max</sub> (mg/g) | a | b (mg/l) | c | R<sup>2</sup> |
| Lang. | 77.2 | 1 | 0.085 | 1 | 0.98600 |
| H-S | 102.9 | 0.487 | 0.218 | 1 | 0.99400 |
| B-S | 103.1 | 0.336 | 0.709 | 0 | 0.99600 |
| B-G | 103 | 0.408 | 0.361 | 0.5 | 0.99550 |
| R-P |  | 0.78 |  |  | 0.99700 |

It is worth to note that using only R^2^ to determine the best-fitting model is not sufficient and could lead to some ambiguities when the set of data is not complete. Indeed, the results in Table 2 showed that based on R^2^ values, H-S, B-G and R-P models seem to be adequate and the parameters of these models were the best fitting for the experiments results. It must be noted however that the R-P model which does not have the proper asymptotic behaviors is generally less performing than the other isotherm here that we have complete data. The calculation of the error deviation using another methods such as the average relative error and/or the Marquardt's percent standard error may be also of interest to describe the validity of sorption model as suggested by **Ncibi et al. (2008)** for the adsorption of phenol and methylene blue (MB), respectively onto a non-porous adsorbent, Posidonia oceanica fibers and two porous adsorbents, chemically and physically activated carbons prepared from vetiver roots.

Several efforts have been made to characterize the metal binding properties of various forms of biomass. The uptake of Pb(II) by different biomass is summarized in Table (3). It can be seen that the algal biomass has a high biosorption capacity, when compared with other biomass, which may be owing to the fact that this biosorbent has many active groups that enhance the biosorption process.

**Table 3.**
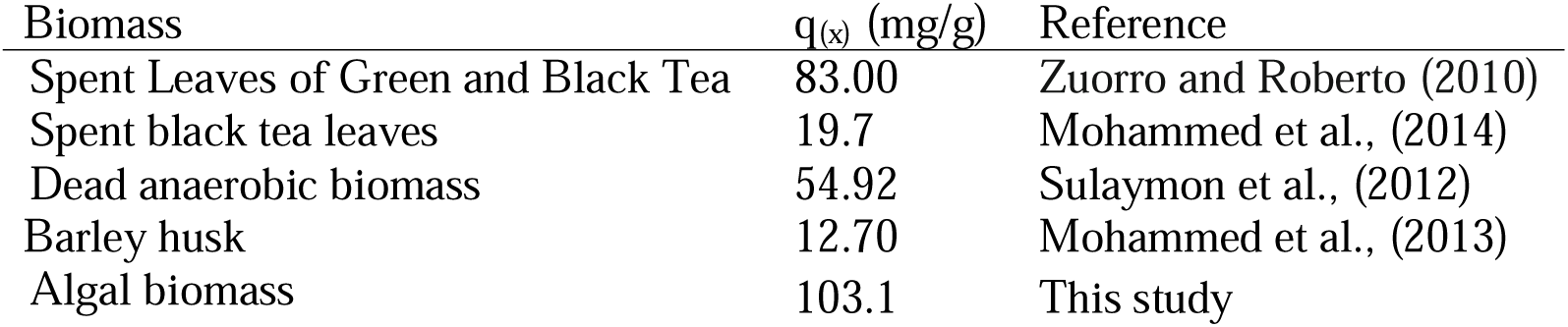
Uptake of Pb(II) by different biomasses

## 4. Conclusions

This study revealed the following conclusions:

- This study confirmed that algal biomass is a promising biosorbent for Pb(II) removal from aqueous solution and the maximum removal efficiency was reach 98% at 40 °C and pH 3.
- Excellent results were obtained by using original non linear isotherm models and one should leave the old-fashion linear methods now that one has highly speed computers and that performing nonlinear recursion methods are on the market.
- The experimental isotherm data were better described by the Hill-Sips and Brouers–Sotolongo model involving the biosorption of Pb(II) on a heterogeneous surface onto algal biomass particles.
- This study shows that a complete set of data is necessary to have a good representation of the isotherm.
- Using only coefficient of determination is not always an appropriate tool to compare the goodness of the non linear fit of an isotherm models.

## Acknowledgments

We would like to express our sincere thanks and deep gratitude to Professor Sarra Gaspard (France) and Professor Mongi Seffen (Tunis) for their useful discussions on the use of the non-linear approach. Special thanks to the Environmental Engineering Department, University of Baghdad, Iraq for supporting this work.

